# Host gene expression changes induced by an HIV-1 mutant associated with delayed AIDS progression reveal potential mechanisms

**DOI:** 10.64898/2026.02.18.706531

**Authors:** Sidney T. Sithole, Joshua Ramsey, Macey C. Call, Brian D. Poole, Brett E. Pickett, Bradford K. Berges

**Affiliations:** Department of Microbiology and Molecular Biology, Brigham Young University, Provo, Utah, USA

**Keywords:** HIV, AIDS, Vpr, apoptosis, RNA sequencing, transcriptomics

## Abstract

HIV-1 viral protein R (Vpr) is a multifunctional protein central to HIV-1 pathogenesis and progression to AIDS. Polymorphisms in *vpr* have been linked to varying rates of HIV-1 progression. Of note is the HIV-1 Vpr R77Q mutant, associated with delayed progression to AIDS (also as the long-term non-progressor phenotype, or LTNP). We previously demonstrated that the R77Q mutant promotes a non-inflammatory, apoptotic phenotype in CD4+ T cells. To investigate the mechanism underlying the R77Q-induced apoptotic phenotype, we performed RNA sequencing on a CD4+ T cell line, HUT78, infected with either a replication-competent wild-type strain (NL4-3) or the R77Q mutant. Our results show that at 72 hours post-infection, transcriptomes were heterogeneous, and differential expression analysis identified 289 differentially expressed genes (DEGs) in the R77Q vs. WT comparison. Functional enrichment analysis revealed enriched pathways associated with apoptosis. Gene ontology (GO) terms and GO connections also revealed an apoptotic signature. Although both viral strains upregulated pro-apoptotic genes, the R77Q mutant failed to upregulate some key anti-apoptotic genes such as *bcl-2*, while WT-infected cells displayed upregulation of those anti-apoptotic genes. Predicted protein-protein interaction within the Bcl-2 family local network also suggests that interactions within this network were substantially affected. Taken together, these findings provide a transcriptomic basis for our previous observations of enhanced apoptosis in R77Q-infected cells and highlight distinct host cell responses that may underlie delayed HIV-1 progression. These results may be useful in identifying new targets to delay AIDS progression.

**Importance:** The R77Q mutant has been a long-standing variant of interest because of its association with long-term non-progressors (LTNPs), yet the mechanisms behind this phenotype remain poorly defined. We previously showed that the R77Q mutant is less cytotoxic, killing fewer cells than WT via a non-inflammatory apoptotic pathway while suppressing inflammatory cytokine release, possibly contributing to the LTNP phenotype. Here, we provide a transcriptomic basis for this phenotype and identify a potential novel loss-of-function mechanism in which R77Q-induced apoptosis arises from a failure to upregulate key anti-apoptotic genes. Defining host responses in the context of delayed HIV-1 progression offers novel insights into cellular networks that could inform future therapeutic strategies beyond the current antiviral approaches.

## Introduction

Human immunodeficiency virus type 1 (HIV-1) is the main causative agent of acquired immunodeficiency syndrome (AIDS) (1–3). AIDS occurs when HIV infects CD4+ helper T cells, killing them and thus inhibiting adaptive immune responses, leaving patients highly susceptible to diseases that would otherwise be detected by the immune system (4–7). Chronic immune activation and inflammation is a hallmark of AIDS development (8–11), and a variety of factors are thought to contribute to this activation (8, 12, 13). Some patients develop AIDS slowly, or not at all, and are commonly referred to as long-term non-progressors (LTNP) (14, 15). While host genetic factors contribute to the development of the LTNP phenotype (16, 17), substantial evidence exists to support HIV-1 genetics playing a significant role (18–23).

The viral protein R (*vpr*) gene in HIV-1 is one such gene where polymorphisms have been found to be associated with the rate of AIDS development (23–25). The R77Q mutation was identified in 2003 as a nonsynonymous single nucleotide polymorphism (SNP) that was detected at significantly higher prevalence in LTNP patients (20, 26). However, the mechanism(s) by which this mutation could cause LTNP are not well understood (22) . Our group previously reported that the R77Q mutation is associated with higher levels of apoptotic cell death in multiple CD4+ T cell lines as well as in cord blood-derived primary CD4+ T cells (27). In contrast, the wild-type NL4-3 strain killed cells through a mechanism involving membrane permeability, which is indicative of necrosis (28, 29). Furthermore, infected cells showed significantly higher levels of G_2_ cell cycle arrest and lower levels of pro-inflammatory cytokine production (TNF and IL-6), as compared to the wild-type strain. Reduced levels of pro-inflammatory cytokines are a potential link between the R77Q mutant and slower AIDS progression (30, 31), since strong inflammatory signals may be connected to increased chronic immune activation (13, 33–35). However, that study did not identify the signals that contribute to the apoptotic and G_2_ arrest phenotypes (27).

In this study, we performed a bulk transcriptomic analysis (Illumina RNA-Seq) of three populations of cells: a) infection with wild-type strain NL4-3, b) infection with the R77Q mutant (a single point mutation difference as compared to NL4-3, which only affects the *vpr*), and c) mock-infected cells. The HUT78 cell line was used, which was the main cell type used in our previous report (27). Multiple time points were analyzed; however, significant differences in the R77Q vs. WT comparison were only noted at 72 hours post-infection (hpi), which became the focus of subsequent analysis. We performed gene ontology (GO) term analysis and Kyoto Encyclopedia of Genes and Genomes (KEGG) analyses to identify genes and cellular pathways, respectively, that differed in the various samples. We found 289 genes to be significantly differentially expressed when comparing R77Q (case) to WT HIV-1(control). Because of our previous findings related to the induction of apoptosis, we focused on analysis of genes and pathways related to this process. We found that *bcl-2*, a key inhibitor of intrinsic apoptosis (32, 33), was upregulated during infection with WT virus, whereas R77Q failed to induce *bcl-2* expression above control levels. Bim, which normally inhibits Bcl-2 (34–36), had gene expression significantly upregulated by R77Q as compared to WT virus. These findings shed insight into how the R77Q mutant could trigger a host response that has lower levels of pro-inflammatory signals as compared to the WT virus and could help to explain how it potentially causes the LTNP phenotype (37).

## Results

### HIV-1 Vpr R77Q-vs. WT-infected cells show distinct and divergent transcriptomic signatures at 72 hpi

To investigate the mechanism underlying HIV-1 Vpr R77Q–induced apoptosis, we analyzed host transcriptomic changes in the human CD4+ T cell line HUT78. We infected cells at a multiplicity of infection (MOI) of 0.3 with replication-competent HIV-1 NL4-3 wild-type (WT) virus or the R77Q mutant in the presence of 10 µg/mL polybrene. In our previous report, we showed no significant differences in efficiency of virus replication between these two strains (27). We treated mock-infected cells with an equivalent concentration of polybrene and an equal volume of virus-free supernatant. All experimental conditions were performed in triplicate.

We then harvested total RNA at 4, 8, 12, 24, and 72 hours post-infection (hpi). RNA quality assessment demonstrated high integrity across all samples, with RNA Integrity Numbers (RIN) exceeding 9.0 for most samples and a minimum RIN of 8.6 when assessed by LC Sciences, a CRO contracted for sequencing (data not shown). We sequenced sample libraries on an Illumina NovaSeq 6000 platform, yielding an average of 21.5 million reads per sample and approximately 64.5 million reads per condition following quality trimming using FastQC.

To determine differential gene expression due to Vpr sequence variation, we compared transcriptomic profiles from R77Q- and WT-infected cells across multiple time points (4, 8, 12, 24, and 72 hpi). Principal component analysis (PCA) revealed no clear separation between R77Q vs. WT infected cells at 4, 8, 12, or 24 hpi (Fig. 1, A1–A5), which suggests minimal overall differences across samples. In contrast, robust segregation of R77Q and WT-infected samples was observed at 72 hpi, with principal component 1 and 2 accounting for 90% of the total variance (Fig. 1, A5), indicating substantial transcriptomic divergence at this time point.

**Fig. 1.**
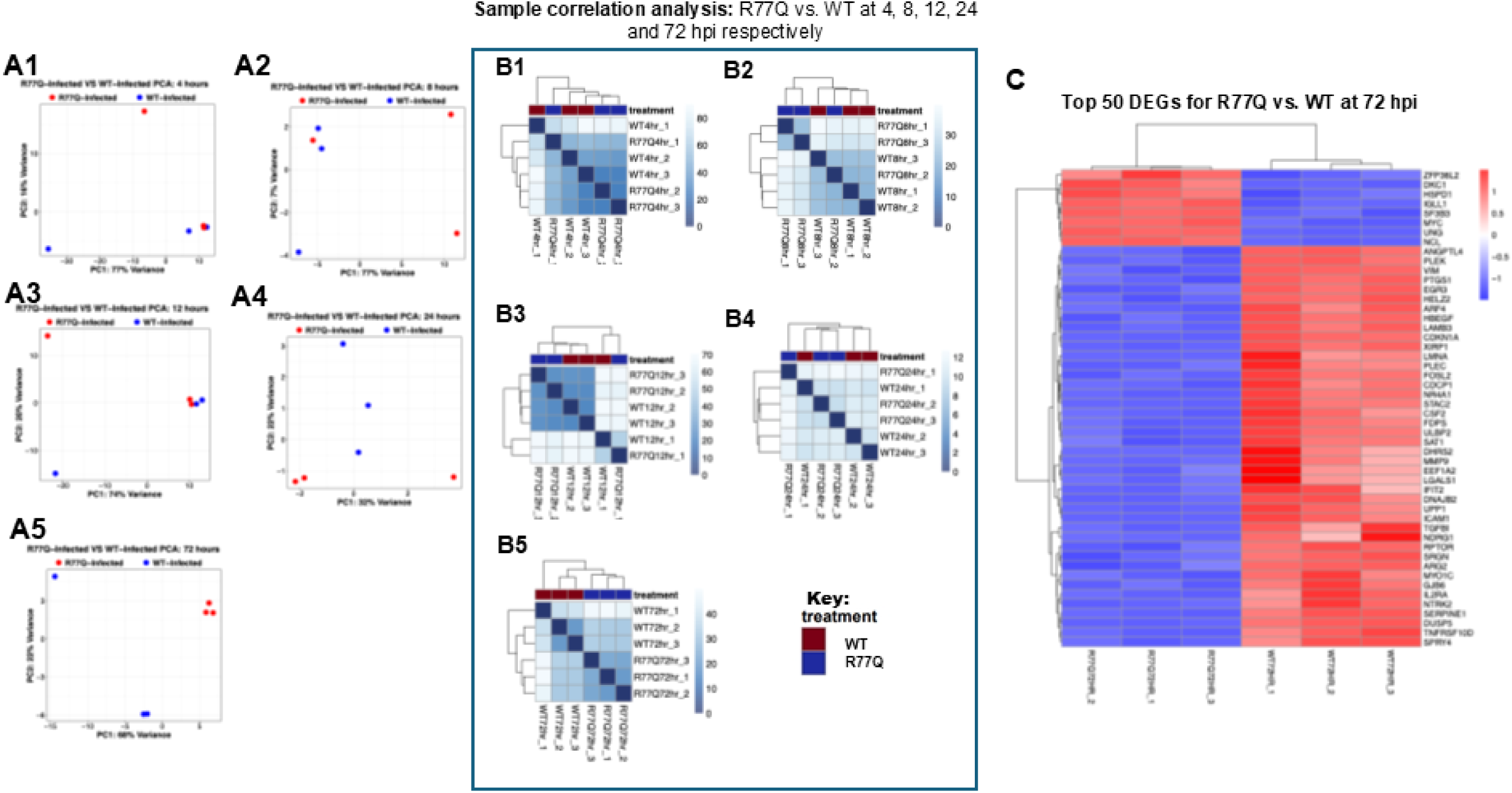
Principal Component Analysis (PCA) and Heatmap analysis show distinct transcriptomic profiles at 72 hpi. (A1-A5) PCA at 4, 8, 12, and 24 hpi, respectively, show no clear separation between conditions. (A5) By 72 hpi, samples cluster distinctly, with the first and second principal components accounting for 68% (PC1) and 22% (PC2) of the variance. (B1-B5) Sample correlation plots at earlier time points show incomplete separation, whereas a clear pattern emerges at 72 hpi (B5). (C) Heatmap of the top 50 significant DEGs at 72hpi scaled by *z-score* demonstrates strong within-group correlation and distinct clustering across conditions.

Sample correlation analysis further supported these findings. Early time points showed limited correlation among replicates (Fig. 1, B1-B4), whereas samples at 72 hpi displayed strong within-group correlation and clear separation between conditions (Fig. 1, B5). Consistent with the PCA and sample distribution analysis, heatmap visualization of the top 50 most significant genes revealed tight clustering of biological replicates within each condition, with highly similar expression patterns indicative of low intra-condition variability. In contrast, samples from different conditions segregated into distinct clusters and exhibited contrasting expression patterns across gene blocks, consistent with condition-specific transcriptional signatures (Fig. 1, C). Based on quality control and variance assessments, all subsequent differential expression and pathway analyses for the R77Q vs. WT comparison were performed using data from the 72 hpi time point.

### Differentially Expressed Genes (DEGs) in R77Q vs. WT infected cells

We determined differentially expressed genes (DEGs) by using a false discovery rate (FDR) adjusted p-value threshold below 0.05. Despite differing by a single nucleotide and amino acid, with genome similarity of about 99.99%, WT and R77Q induced 289 DEGs in direct comparison when applying a filtering threshold of |log_2_fold change (logFC)| ≥ 1 (i.e logFC ≤ -1 or ≥ 1). Relative to mock infection, WT infection resulted in 2014 upregulated and 1624 downregulated genes (Fig. 2, A1), whereas R77Q infection led to 1574 upregulated and 1362 downregulated genes (Fig. 2, A2). Direct comparison of R77Q vs. WT revealed 238 upregulated and 51 downregulated genes (Fig. 2, A3).

**Fig. 2.**
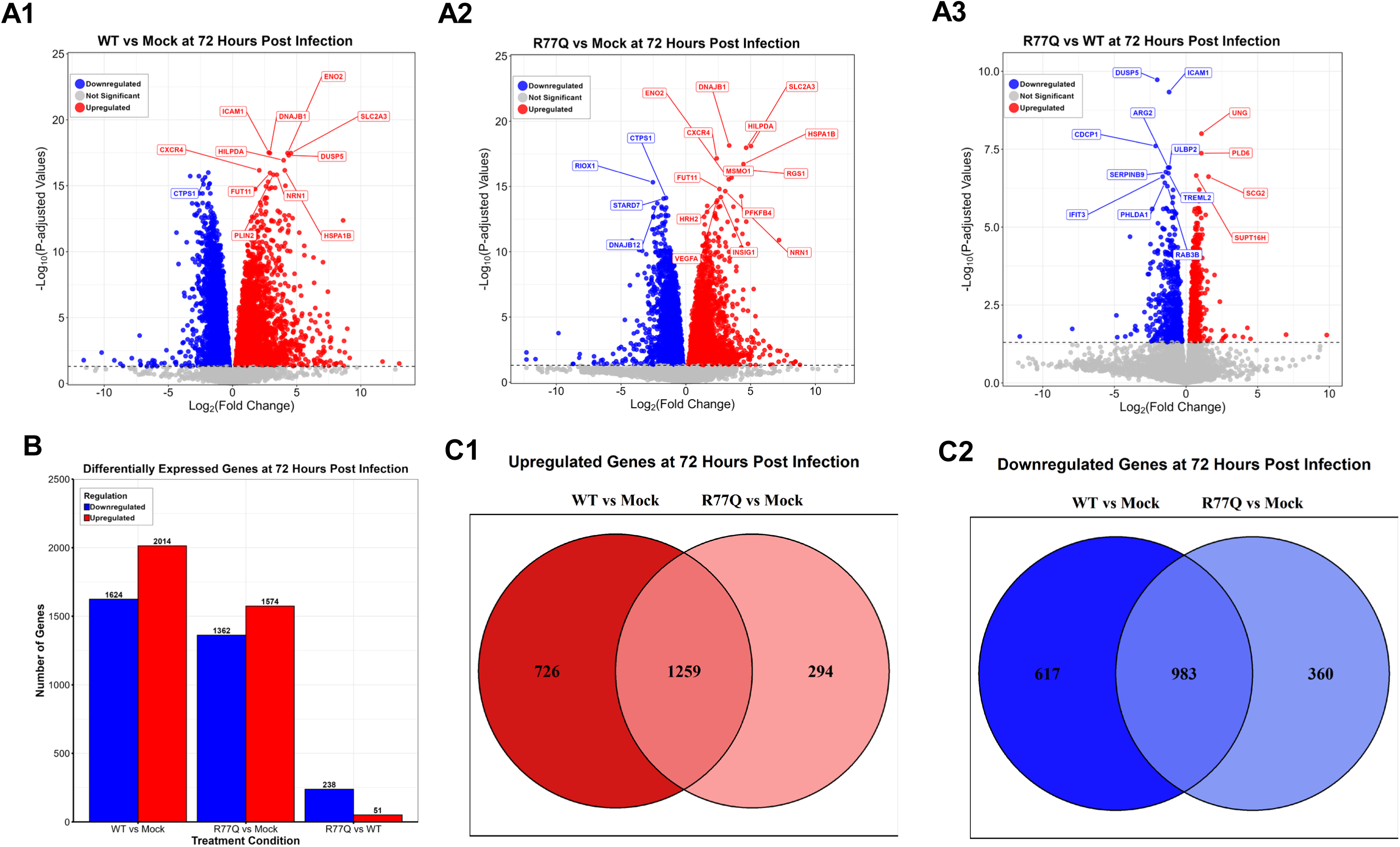
Differential gene expression (DGE) analysis at 72 hpi. A total of 289 differentially expressed genes (DEGs) were observed in R77Q vs. WT-infected cells 72hpi. (A1-A3) Volcano plots of WT vs. mock, R77Q vs. mock, and R77Q vs. WT, respectively. Red and blue points represent upregulated and downregulated DEGs respectively while grey dots represent non-significant genes (FDR <0.05) (B) Bar graph summarizing the number of DEGs for each comparison at 72 hpi. (C1-C2) Venn diagrams showing the overlap of DEGs, separated into downregulated (C1) and upregulated (C2) sets for each condition.

We observed substantial overlap when comparing transcriptional differences between cells infected with WT and R77Q viruses. When compared to mock infection, 983 genes were commonly downregulated in both conditions (Fig. 2, C1), and 1259 genes were commonly upregulated (Fig. 2, C2), indicating a largely similar host response with quantitative difference between variants. Furthermore, after ranking the top 10 DEGs by highest FDR in the R77Q vs. WT comparison, we identified a subset of highly regulated genes, the majority of which have established roles in apoptosis, inflammation, DNA damage response, or cytokine signaling (Supplementary Table S1).

### Functional Enrichment Analysis reveals an enriched apoptotic pathway

To define biological processes and pathways underlying the transcriptional differences induced by infection with WT virus and the R77Q mutant, we performed GO enrichment and KEGG pathway enrichment analyses using the gprofiler2 R package, version v0.2.4. After filtering GO terms for apoptosis-related terms, analysis of upregulated genes revealed enrichment of apoptosis-associated GO terms in both WT vs. mock and R77Q vs. mock comparisons, indicating activation of apoptotic transcriptional programs by both Vpr sequences. Notably, WT vs. mock infection was additionally enriched for GO terms related to negative regulation of apoptosis, a feature not observed in the R77Q vs. mock comparison (Fig. 3) (Table 1).

**Fig. 3.**
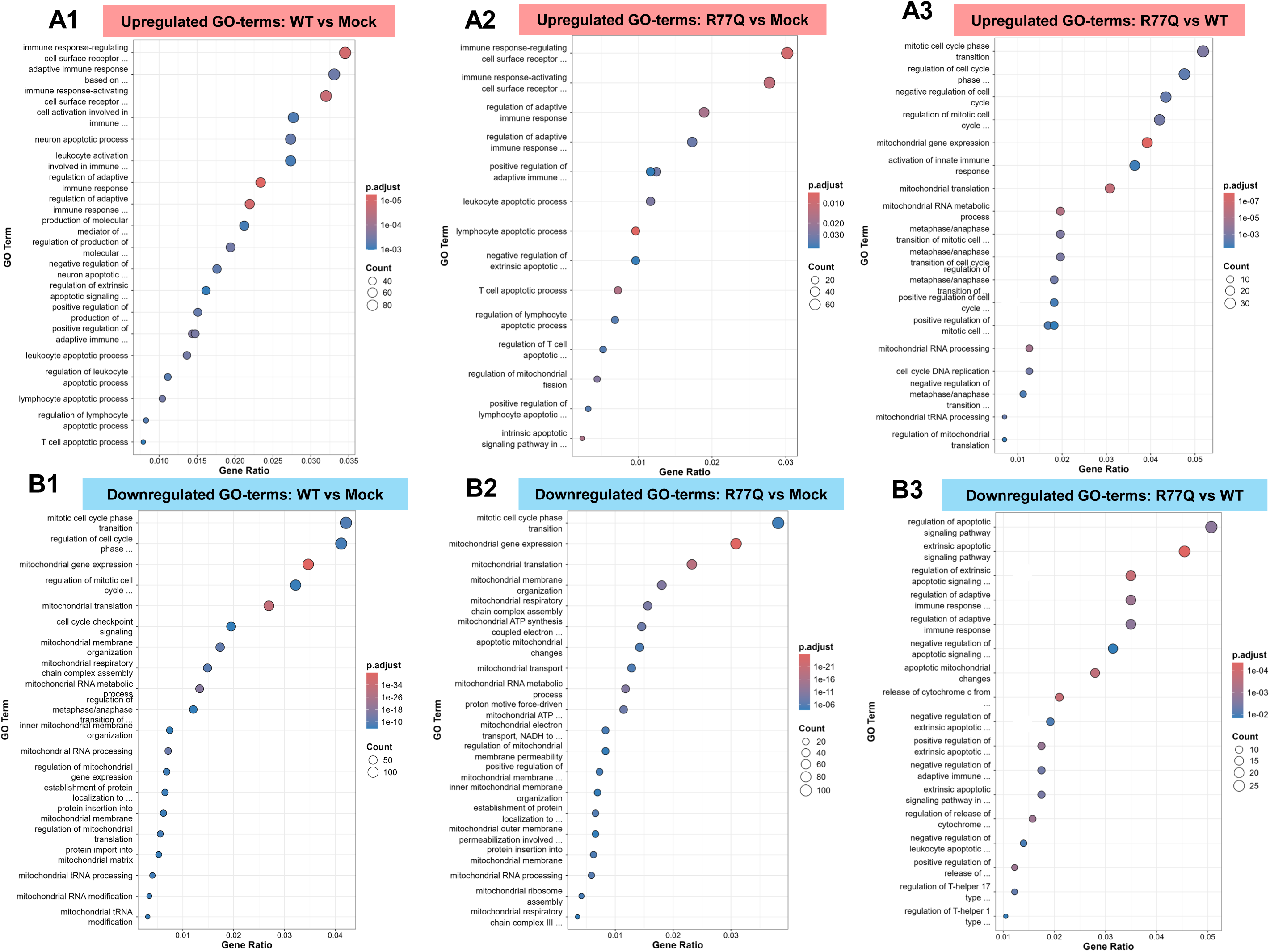
Gene ontology (GO) analysis of Biological Processes (BP) at 72hpi highlight apoptosis-related terms. (A1-A3) Dot plot showing the top significantly upregulated apoptosis-related GO terms for WT vs. Mock, R77Q vs. Mock, and R77Q vs. WT respectively. (B1-B3) Dot plot showing the top significantly downregulated apoptosis-related GO terms for WT vs. Mock, R77Q vs. Mock, and R77Q vs. WT respectively

**Table 1:** Apoptosis-related GO: BP terms significantly upregulated in the WT vs. Mock comparison.

| GO: BP Term Description | Gene Ratio | Fold Enrichment | zScore | p.adjust |
| --- | --- | --- | --- | --- |
| Regulation of adaptive immune response | 65/2782 | 1.967206 | 6.057433 | 5.74E-06 |
| Regulation of adaptive immune response based on somatic recombination of immune receptors built from immunoglobulin superfamily domains | 61/2782 | 1.969224 | 5.875113 | 1.05E-05 |
| Immune response-regulating cell surface receptor signaling pathway | 96/2782 | 1.690422 | 5.693436 | 1.12E-05 |
| Immune response-activating cell surface receptor signaling pathway | 89/2782 | 1.709227 | 5.595344 | 1.74E-05 |
| Positive regulation of adaptive immune response based on somatic recombination of immune receptors built from immunoglobulin superfamily domains | 40/2782 | 2.054332 | 5.056389 | 2.27E-04 |
| Positive regulation of adaptive immune response | 41/2782 | 2.014139 | 4.973734 | 2.80E-04 |
| Leukocyte apoptotic process | 38/2782 | 2.060906 | 4.950247 | 3.24E-04 |
| Regulation of production of molecular mediator of immune response | 54/2782 | 1.812287 | 4.82808 | 3.28E-04 |
| Lymphocyte apoptotic process | 29/2782 | 2.312936 | 5.046438 | 3.28E-04 |
| Adaptive immune response based on somatic recombination of immune receptors built from immunoglobulin superfamily domains | 92/2782 | 1.551481 | 4.649142 | 3.82E-04 |
| Neuron apoptotic process | 76/2782 | 1.620209 | 4.639757 | 4.46E-04 |
| Positive regulation of production of molecular mediator of immune response | 42/2782 | 1.950208 | 4.794395 | 4.51E-04 |
| Negative regulation of neuron apoptotic process | 49/2782 | 1.835279 | 4.696949 | 5.19E-04 |
| Regulation of leukocyte apoptotic process | 31/2782 | 2.166579 | 4.791492 | 6.10E-04 |
| Regulation of lymphocyte apoptotic process | 23/2782 | 2.474981 | 4.877916 | 6.43E-04 |
| Cell activation involved in immune response | 77/2782 | 1.586644 | 4.465239 | 7.05E-04 |
| Leukocyte activation involved in immune response | 76/2782 | 1.585312 | 4.427462 | 7.96E-04 |
| Production of molecular mediator of immune response | 59/2782 | 1.694824 | 4.468031 | 8.31E-04 |
| Regulation of extrinsic apoptotic signaling pathway | 45/2782 | 1.826756 | 4.463938 | 0.001008 |
| T cell apoptotic process | 22/2782 | 2.444992 | 4.702024 | 0.001036 |
| Negative regulation of immune response | 56/2782 | 1.679825 | 4.276815 | 0.001427 |
| Lymphocyte activation involved in immune response | 57/2782 | 1.64434 | 4.134496 | 0.002033 |
| Cytokine production involved in immune response | 36/2782 | 1.877343 | 4.175363 | 0.002434 |
| Regulation of cytokine production involved in immune response | 36/2782 | 1.877343 | 4.175363 | 0.002434 |
| Regulation of neuron apoptotic process | 62/2782 | 1.592107 | 4.030087 | 0.002529 |
| Positive regulation of cytokine production involved in immune response | 27/2782 | 2.080011 | 4.224119 | 0.002662 |
| Negative regulation of extrinsic apoptotic signaling pathway | 30/2782 | 1.974552 | 4.125379 | 0.00309 |
| Regulation of T cell apoptotic process | 16/2782 | 2.522529 | 4.157635 | 0.0046 |
| Mature B cell differentiation involved in immune response | 13/2782 | 2.592084 | 3.864988 | 0.01018 |
| Extrinsic apoptotic signaling pathway | 54/2782 | 1.531724 | 3.441164 | 0.011075 |
| Positive regulation of innate immune response | 88/2782 | 1.380968 | 3.332059 | 0.012407 |
| Regulation of immunoglobulin mediated immune response | 19/2782 | 2.077526 | 3.535002 | 0.014979 |
| Positive regulation of lymphocyte apoptotic process | 9/2782 | 3.050683 | 3.816761 | 0.015688 |
| Intrinsic apoptotic signaling pathway in response to endoplasmic reticulum stress | 20/2782 | 2.02367 | 3.491575 | 0.01597 |
| Negative regulation of leukocyte apoptotic process | 18/2782 | 2.103919 | 3.502444 | 0.016445 |
| Intrinsic apoptotic signaling pathway in response to hypoxia | 6/2782 | 4.067577 | 4.036098 | 0.016445 |
| CD4-positive, alpha-beta T cell differentiation involved in immune response | 25/2782 | 1.8422 | 3.368387 | 0.017972 |
| Innate immune response-activating signaling pathway | 71/2782 | 1.407398 | 3.162688 | 0.019054 |
| Regulation of apoptotic signaling pathway | 82/2782 | 1.369217 | 3.128396 | 0.019853 |
| Alpha-beta T cell activation involved in immune response | 25/2782 | 1.822391 | 3.307078 | 0.020192 |
| Alpha-beta T cell differentiation involved in immune response | 25/2782 | 1.822391 | 3.307078 | 0.020192 |
| Negative regulation of adaptive immune response | 21/2782 | 1.923854 | 3.312257 | 0.021593 |
| Antiviral innate immune response | 22/2782 | 1.887905 | 3.289596 | 0.021965 |
| Negative regulation of adaptive immune response based on somatic recombination of immune receptors built from immunoglobulin superfamily domains | 20/2782 | 1.936942 | 3.266781 | 0.023982 |
| Negative regulation of lymphocyte apoptotic process | 13/2782 | 2.319233 | 3.386119 | 0.024608 |
| Negative regulation of apoptotic signaling pathway | 53/2782 | 1.466541 | 3.05748 | 0.025439 |
| T cell differentiation involved in immune response | 26/2782 | 1.745165 | 3.123419 | 0.027899 |
| Activation of innate immune response | 73/2782 | 1.367095 | 2.933546 | 0.030566 |
| Regulation of extrinsic apoptotic signaling pathway via death domain receptors | 15/2782 | 2.075295 | 3.135028 | 0.035785 |
| Immunoglobulin production involved in immunoglobulin-mediated immune response | 17/2782 | 1.953356 | 3.050792 | 0.038037 |
| Type 2 immune response | 14/2782 | 2.109114 | 3.098505 | 0.038159 |
| Regulation of endoplasmic reticulum stress-induced intrinsic apoptotic signaling pathway | 12/2782 | 2.259765 | 3.147073 | 0.038159 |
| Positive regulation of cell cycle G2/M phase transition | 11/2782 | 2.330383 | 3.133079 | 0.040452 |
| T cell activation involved in immune response | 33/2782 | 1.57547 | 2.863249 | 0.04119 |
| Somatic recombination of immunoglobulin genes involved in immune response | 16/2782 | 1.972159 | 3.003335 | 0.041502 |
| Somatic diversification of immunoglobulins involved in immune response | 16/2782 | 1.972159 | 3.003335 | 0.041502 |
| Positive regulation of T cell apoptotic process | 7/2782 | 2.965942 | 3.272397 | 0.043788 |
| Intrinsic apoptotic signaling pathway | 67/2782 | 1.355859 | 2.733652 | 0.044599 |
| T cell mediated immune response to tumor cell | 8/2782 | 2.711718 | 3.185875 | 0.045065 |
| Positive regulation of leukocyte apoptotic process | 11/2782 | 2.259765 | 3.012853 | 0.049204 |

**Table 2:** Apoptosis-related GO:BP terms significantly upregulated in the R77Q vs. Mock comparison.

| Description | Gene Ratio | Fold Enrichment | zScore | p.adjust |
| --- | --- | --- | --- | --- |
| Lymphocyte apoptotic process | 24/2484 | 2.143791 | 4.116208 | 0.006813 |
| Immune response-regulating cell surface receptor signaling pathway | 75/2484 | 1.479076 | 3.698914 | 0.008935 |
| Immune response-activating cell surface receptor signaling pathway | 69/2484 | 1.484105 | 3.575935 | 0.011828 |
| T cell apoptotic process | 18/2484 | 2.240437 | 3.779232 | 0.015124 |
| Intrinsic apoptotic signaling pathway in response to hypoxia | 6/2484 | 4.555556 | 4.38009 | 0.016012 |
| Regulation of adaptive immune response | 47/2484 | 1.593089 | 3.477751 | 0.017012 |
| Regulation of mitochondrial fission | 11/2484 | 2.694146 | 3.676624 | 0.026315 |
| Leukocyte apoptotic process | 29/2484 | 1.761481 | 3.326739 | 0.028182 |
| Positive regulation of adaptive immune response | 31/2484 | 1.705582 | 3.23998 | 0.03156 |
| Regulation of adaptive immune response based on somatic recombination of immune receptors built from immunoglobulin superfamily domains | 43/2484 | 1.554674 | 3.148031 | 0.033578 |
| Positive regulation of lymphocyte apoptotic process | 8/2484 | 3.037037 | 3.549807 | 0.039571 |
| Regulation of lymphocyte apoptotic process | 17/2484 | 2.048795 | 3.247489 | 0.039571 |
| Regulation of T cell apoptotic process | 13/2484 | 2.295435 | 3.312118 | 0.04009 |
| Negative regulation of extrinsic apoptotic signaling pathway | 24/2484 | 1.769148 | 3.048442 | 0.047009 |
| Positive regulation of adaptive immune response based on somatic recombination of immune receptors built from immunoglobulin superfamily domains | 29/2484 | 1.66807 | 2.999813 | 0.04789 |

Analysis of downregulated GO terms showed shared enrichment for mitochondrial gene expression and cell cycle–related processes in both WT and R77Q infections compared to mock. Direct comparison of R77Q vs. WT revealed upregulation of GO terms associated with intrinsic apoptosis whereas downregulated GO terms were predominantly linked to regulation of extrinsic apoptotic signaling.

Our GO gene–concept network analysis of the R77Q vs. WT comparison further resolved functional differences. Among the top 10 enriched themes in the R77Q vs. WT comparison, genes involved in the response to virus were largely downregulated, including multiple antiretroviral restriction factors such as APOBEC and TRIM family members, although select genes involved in viral innate immune response such as IFI16 and RIG1were upregulated. In contrast, RNA splicing and regulation of DNA metabolic processes were enriched and predominantly upregulated (Fig. 4).

**Fig. 4.**
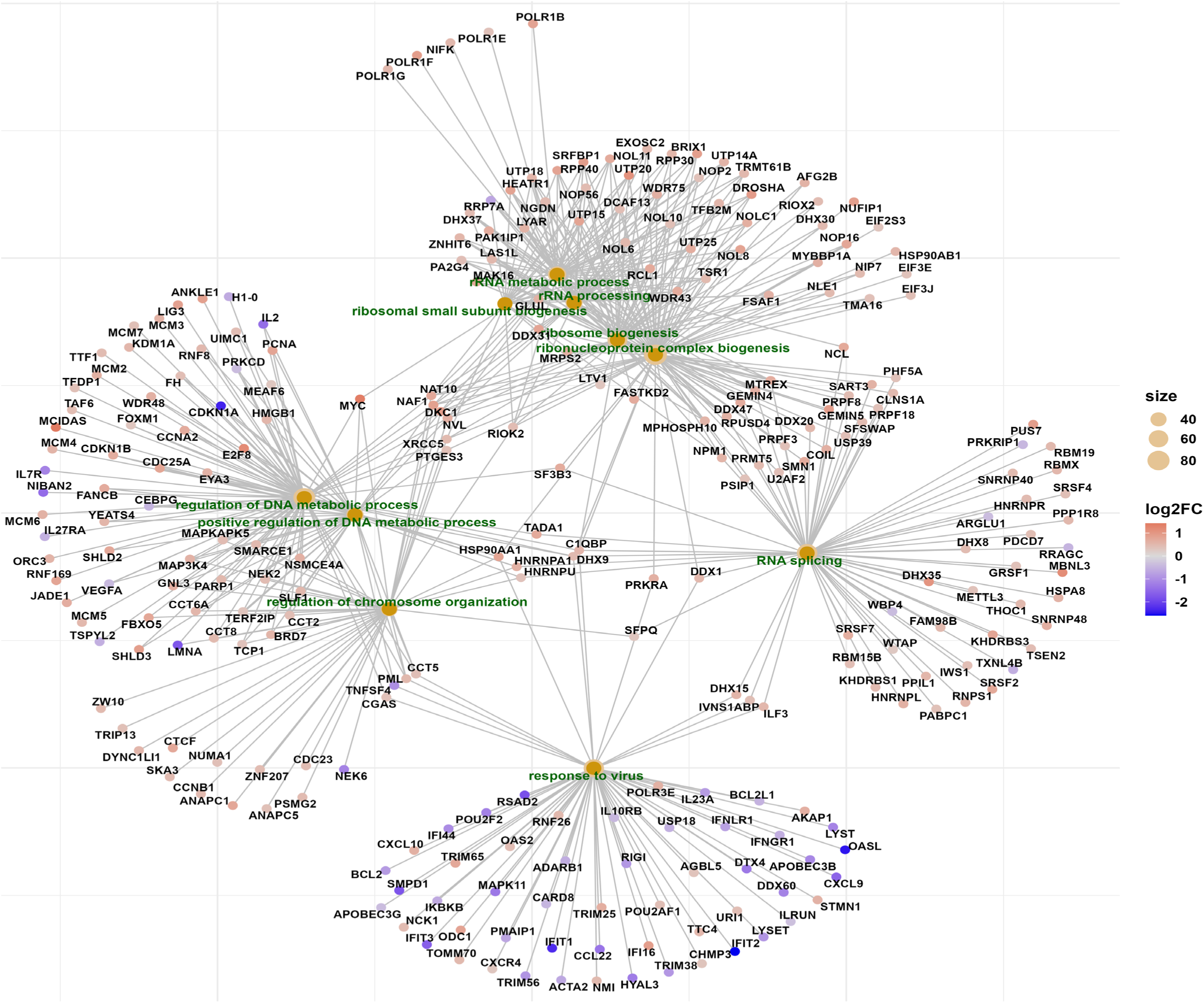
Gene ontology concept network (CNET) of enriched biological processes in the R77Q vs. WT comparison. The CNET plot was generated from significant DEGs (FDR <0.05) in R77Q vs. WT infection at 72 hpi using clusterProfiler. The enriched Gene Ontology Biological Processes are denoted by orange nodes, with node size proportional to the number of genes, as indicated by the scale. Additionally, the enriched GO terms are highlighted in green for clarity. Individual gene nodes are colored according to direction of differential expression in R77Q relative to WT, with red indicating upregulated genes and blue indicating downregulated genes. Edges represent gene membership within each functional category.

Consistent with these findings, KEGG pathway analysis of the R77Q vs. WT comparison identified downregulation of JAK–STAT, TNF signaling, and apoptotic pathways, while pathways related to cell cycle progression and regulation of DNA metabolic processes were upregulated (Fig. S1). Across all enriched pathways, apoptosis also emerged as a recurrent and significantly enriched process, highlighting apoptotic signaling as an important feature (Fig. 5).

**Fig. 5.**
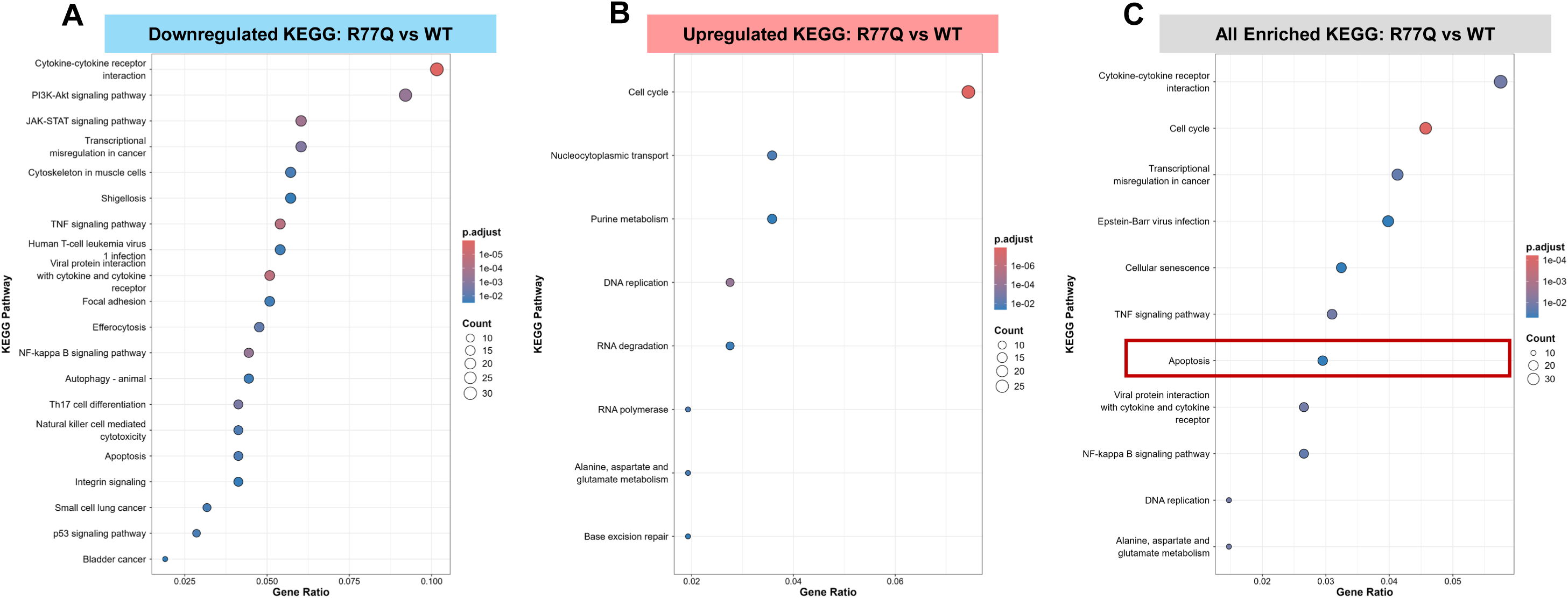
KEGG pathway enrichment analysis of differentially expressed genes in R77Q vs. WT at 72 hpi. KEGG pathway enrichment analysis was performed using gprofiler2 (v0.2.4) on significantly differentially expressed genes (FDR <0.05) identified in the direct comparison between R77Q and WT infection at 72 hpi. Dot plots display enriched KEGG pathways for (A) genes downregulated in the R77Q condition relative to WT, (B) genes upregulated in R77Q vs. WT, and (C) all differentially expressed genes irrespective of direction. Dot size is proportional to the number of genes contributing to each pathway, and dot color reflects the level of statistical significance, as indicated by the scale. The apoptosis pathway is highlighted with a red bounding box and was selected for downstream visualization using Pathview to map differential gene expression onto canonical KEGG apoptosis pathway.

### R77Q is unable to prevent apoptotic signals at the mitochondrial membrane

To further define how WT and the R77Q mutant differentially regulate apoptotic signaling, we interrogated the enriched apoptosis pathway using Pathview to overlay differential gene expression onto the KEGG apoptosis map. This analysis revealed coordinated changes in both pro-apoptotic and pro-survival gene expression in the direct R77Q vs. WT comparison (Fig. 6)

**Fig. 6.**
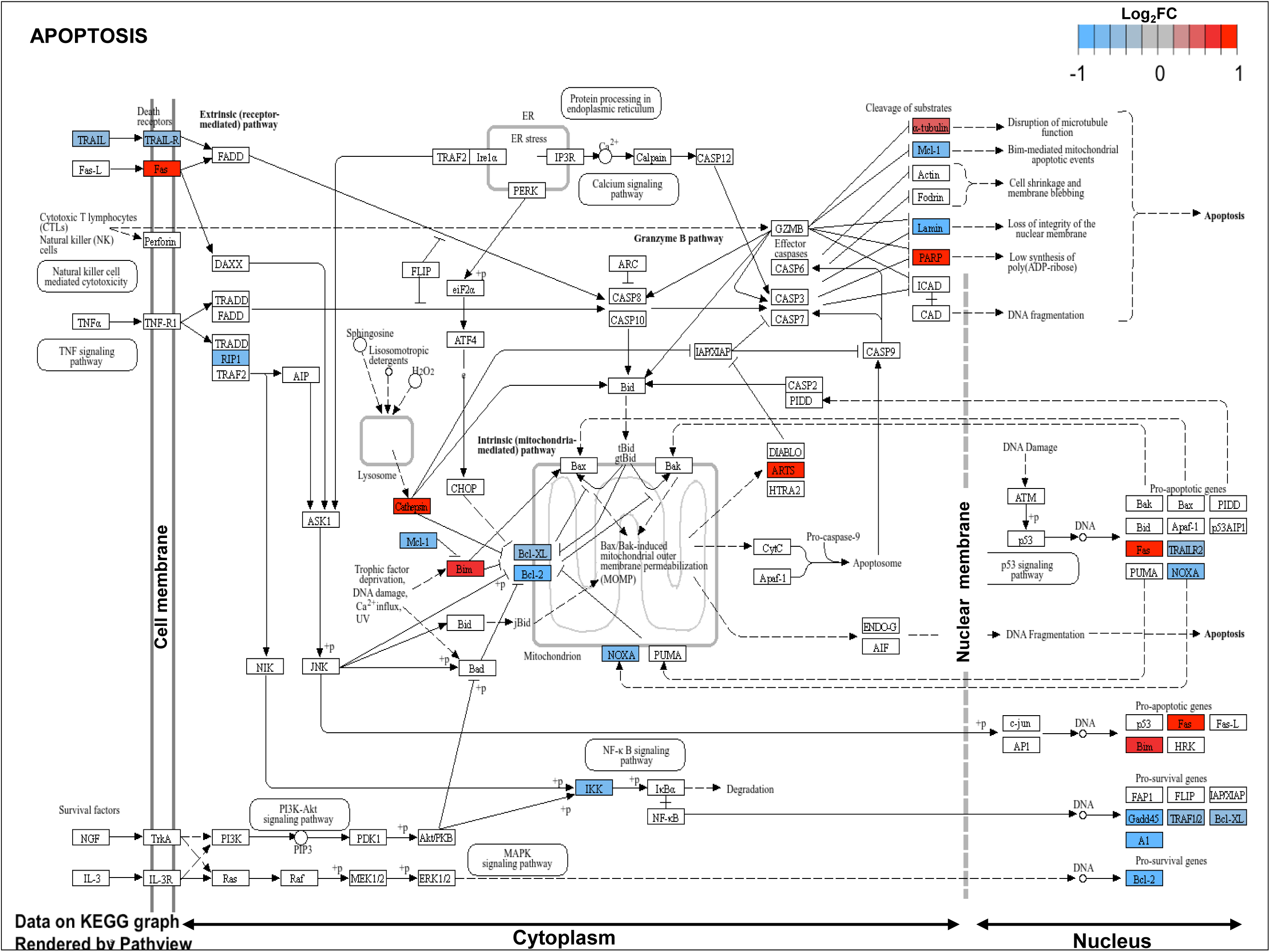
KEGG pathway analysis shows DEGs in the enriched apoptosis pathway in R77Q vs. WT at 72 hpi. The KEGG apoptosis pathway was rendered using Pathview, with differential gene expression data from the R77Q vs. WT comparison at 72 hpi superimposed onto the pathway map. Genes significantly upregulated in the R77Q condition relative to WT are shown in red, whereas genes significantly downregulated are shown in blue. Uncolored pathway components indicate genes that were not differentially expressed in this analysis. This visualization highlights coordinated alterations in apoptotic signaling associated with R77Q infection.

**Fig. 7.**
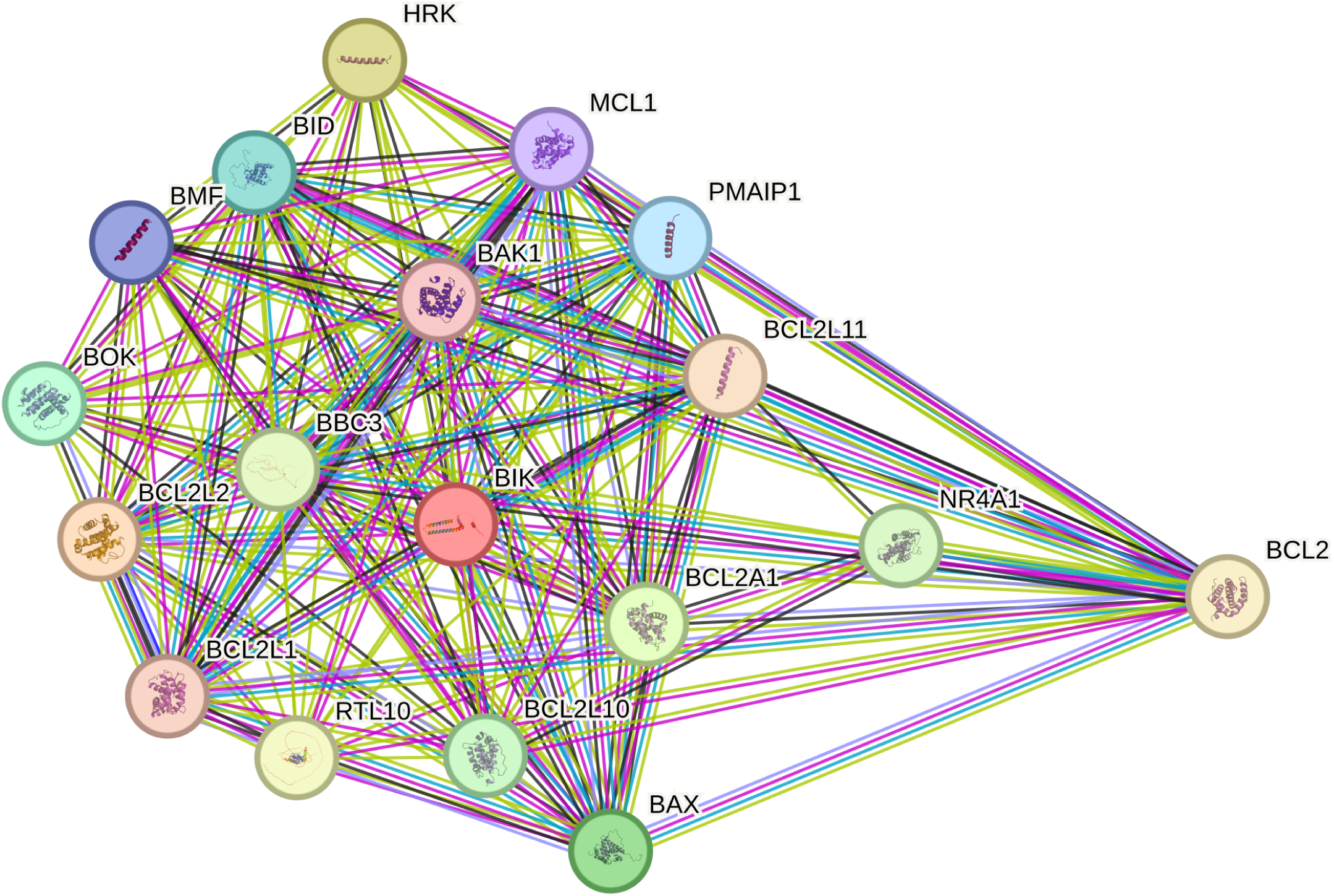
STRING protein-protein interaction analysis of the Bcl-2 family local network cluster in R77Q vs. WT at 72hp. The protein-protein interaction (PPI) analysis was performed using STRING to examine interactions among differentially regulated Bcl-2 family members. Each node represents an individual protein, and edges represent known or predicted physical or functional interactions between proteins. The local network cluster is comprised of 18 interacting proteins (nodes), compared with proteins expected by chance, indicating substantial enrichment. The average node degree of 15.2 reflects the high number of interactions per protein within the cluster, while the average local clustering coefficient of 0.96 indicates that proteins are not only highly connected to a central network but also strongly interconnected with one another. Consistent with this, the PPI enrichment test was significant (p < 1 × 10^-16^), supporting a nonrandom, functionally coherent interaction network among differentially regulated Bcl-2 family members.

Within the R77Q vs. WT comparison, we observed differentially expressed genes associated with extrinsic apoptotic signaling. *FAS*, which encodes the death receptor Fas that initiates extrinsic apoptosis upon ligand engagement, was significantly upregulated in R77Q-infected cells relative to WT (logFC = 1.13; FDR = 9.78 × 10⁻⁵). In contrast, *TNFSF10* (*TRAIL*), encoding a pro-apoptotic death ligand, and its receptor *TNFRSF10* were downregulated (logFC = −0.57; FDR = 0.00226 and logFC = −0.551; FDR = 0.00688, respectively). Despite these changes at the receptor–ligand level, downstream components required to propagate the extrinsic apoptotic cascade were not enriched, consistent with GO analyses indicating overall suppression of extrinsic apoptotic signaling in the R77Q vs. WT comparison (Fig. 3).

In contrast, pronounced differences were observed in the regulation of intrinsic apoptosis–associated genes between WT and R77Q infection. Several anti-apoptotic genes of the Bcl-2 family were downregulated, while pro-apoptotic regulators were enriched. Notably, the anti-apoptotic gene *bcl-2*, which encodes the Bcl-2 protein that inhibits ANT1-mediated mitochondrial pore formation and cytochrome c release (38–40), was significantly upregulated in WT-infected cells (logFC = 1.03; FDR = 7.39 × 10⁻⁴) but not in R77Q-infected cells (logFC = 0.20; FDR = 0.517). Direct comparison confirmed significantly reduced *bcl-2* expression in R77Q relative to WT infection (logFC = −0.83; FDR = 0.0225). Conversely, *BCL2L11* (*bim*), a pro-apoptotic gene that encodes a BH3-only protein (Bim) that is activated in response to DNA damage and functions as a potent antagonist of Bcl-2–mediated survival signaling, was significantly upregulated in R77Q-infected cells relative to WT (logFC = 0.709; FDR = 0.0256).

Taken together, these data indicate a coordinated downregulation of anti-apoptotic gene expression at the mitochondrial membrane in R77Q-infected cells. In contrast to WT, which retains the ability to induce *bcl-2* expression and reinforce mitochondrial survival signaling, the R77Q mutant fails to mount this anti-apoptotic response, consistent with a loss-of-function phenotype in the regulation of mitochondrial apoptotic pathways by the R77Q mutant relative to the WT virus.

### Protein–protein interaction analysis identifies Bcl-2 family–associated networks

To further assess whether the transcriptional differences observed between the R77Q mutant and the WT virus converge on coordinated protein networks, we performed protein–protein interaction (PPI) analysis using the Search Tool for the Retrieval of Interacting Genes/Proteins (STRING) database on differentially expressed genes (FDR < 0.05) from the R77Q vs. WT comparison at 72 hpi. This analysis revealed a highly interconnected interaction network, consisting of 18 nodes and 137 edges. Nodes represent individual proteins and edges represent known or predicted functional interactions. The resulting 137 edges substantially exceeds the expected number of interactions for a random gene set of comparable size (expected edges = 3). The resulting network exhibited a high average node degree (15.2), which represents the average number of interaction partners associated with a given protein in the network with a strong tendency toward clustering (average local clustering coefficient = 0.96), indicating extensive functional connectivity among these proteins. Consistent with this structure, the STRING PPI enrichment analysis was highly significant (p < 1 × 10⁻¹⁶), supporting the conclusion that these proteins form a biologically coherent interaction network rather than a random assembly.

Notably, the interaction network was dominated by proteins associated with mitochondrial apoptosis and Bcl-2 family–mediated regulation of cell death, consistent with the transcriptional enrichment of apoptotic pathways observed in GO and KEGG analyses. The resulting network was centered on regulators of mitochondrial apoptosis, with hub nodes including Bcl-2 family members and associated apoptotic effectors. Prominent hubs included the anti-apoptotic proteins Bcl-2, Bcl-xL (BCL2L1), Bcl-w (BCL2L2), and Mcl-1, alongside pro-apoptotic factors Bax, Bim (BCL2L11), Bid, Bmf, Bok, Hrk, Puma (BBC3), and Noxa (PMAIP1). In addition, NR4A1, a nuclear receptor known to modulate Bcl-2–dependent apoptotic signaling through mitochondrial translocation (41), was identified as a significantly connected node. The prominence of both pro- and anti-apoptotic regulators within this network highlights extensive crosstalk at the mitochondrial membrane. Several highly connected nodes corresponded to key regulators of mitochondrial membrane integrity and apoptotic signaling, reinforcing the notion that differential regulation of apoptosis-related genes in the presence of the R77Q mutant reflects coordinated disruption of protein interaction networks rather than isolated transcriptional changes.

## Discussion

HIV-1 Vpr polymorphisms are linked to differential rates of HIV-1 disease progression, highlighting how subtle viral sequence variations can translate into meaningful biological outcomes (18, 42–44). Among these, the Vpr R77Q mutation has been repeatedly associated with delayed progression to AIDS, yet the mechanistic basis for this phenotype has remained unclear (45). Earlier studies proposed reduced cell death in R77Q infection as the sole basis for the LTNP phenotype (45–48); however, subsequent work by our group demonstrated that this difference is not just solely quantitative but also qualitative, with R77Q preferentially inducing apoptotic rather than necrotic forms of cell death (27). This distinction is biologically significant (49, 50), as apoptosis is comparatively non-inflammatory (51–53), whereas necrotic cell death promotes chronic inflammation (50, 54–56), which is a hallmark of HIV-1 disease progression (11, 57). Despite these observations, the molecular mechanisms linking R77Q-mediated modulation of cell death to alter disease progression remains largely unresolved (54). In this study, we sought to delineate the host transcriptional mechanism underlying R77Q-mediated modulation of apoptotic signaling.

Using transcriptomic profiling, we identified several key features of the host response to R77Q vs. WT infection. First, both WT and R77Q triggered clear transcriptional changes relative to mock-infected cells as early as 4 hpi, yet meaningful differences between the two strains did not emerge until 72 hpi, at which point their host responses began to diverge. Second, nearly 300 DEGs distinguished R77Q from WT infection at this time point. Third, functional enrichment analysis identified that apoptosis and apoptosis-related GO-terms were enriched in both R77Q and WT infections; however, WT infection was uniquely associated with downregulation of intrinsic apoptotic GO-terms, a pattern not shown in R77Q infection. Fourth, KEGG analysis revealed significant enrichment of apoptotic signaling in the R77Q vs. WT comparison and, as visualized by Pathview (58), highlighted a failure of R77Q infection to upregulate key anti-apoptotic regulators such as *bcl-2*, consistent with a loss of function phenotype. Finally, protein-protein interaction analysis revealed a highly significant interaction network centered on the Bcl-2 family of proteins, supporting coordinated regulation of apoptotic signaling involving this network.

During the early phase of infection (4, 8, 12, 24 hpi), both R77Q and WT-infected cells showed clear transcriptomic changes relative to mock-infected cells, consistent with previous work by others (55, 59). Bauby et al. (2021) demonstrated that HIV-1 infection triggers rapid and widespread transcriptomic changes in CD4+ T cells as early as 4–12 hpi, with Vpr driving much of this early host response (59). Similarly, Mohammadi et al, reported broad and rapid host gene expression changes following HIV-1 infection, underscoring the magnitude of early virus-driven transcriptional disruptions (55). Notably, early time points are characterized by extensive differential expression when compared to uninfected controls (60). In contrast, direct comparison between closely related viral strains may reveal far less pronounced differences during the early infection (61). Taken together, these observations suggest that the early transcriptional landscape is dominated by a broad, generalized host response to infection, whereas strain-specific effects become more discernible at later stages (62).

In our model, no significant DEGs were detected when directly comparing R77Q vs. WT at early point. This was expected given the nearly identical viral genomes (∼99.99% identity) and the use of a relatively low multiplicity of infection (MOI = 0.3) in this model, conditions under which strain-specific effects often emerge more slowly (21, 63). Despite the subtlety of the R77Q mutation, the magnitude of transcriptional changes observed at 72 hpi was striking, with nearly 300 DEGs distinguishing R77Q from WT infection. This finding aligns with prior work demonstrating that the *vpr* gene can exert significant effects on host gene expression (64, 65). Consistent with this, Imbeault et al. (2012) showed that small viral mutations can yield only minor transcriptomic differences compared to WT infection. In their experiment model, at 72 hpi, HIV-1 infection generated more than 3,300 DEGs relative to mock, yet one of the Vpu point mutant differed from WT by only about 7 DEGs at the same time point, a pattern expected given how similar the viral strains were (66).

Building on these global transcriptional differences, a closer inspection of the DEGs revealed a consistent enrichment of apoptosis-related genes, in line with the established role of Vpr as a regulator of programmed cell death (23, 67). Within the extrinsic apoptotic pathway, R77Q-infected cells showed selective modulation of death receptor signaling. *FAS*, which encodes the Fas death receptor responsible for initiating the extrinsic apoptosis pathway upon FasL engagement (68–70), was significantly upregulated in R77Q vs. WT-infected cells. However, *FasL* was downregulated, suggesting that, at least at gene level, although receptor expression increased, the corresponding ligand signal required to trigger the pathway was reduced. This imbalance suggests that the extrinsic pathway was primed but not activated (71).

A similar pattern appeared in the TRAIL pathway. *TNFSF10*, which encodes the pro-apoptotic ligand TRAIL, and *TNFRSF10*, which encodes its cognate death receptors DR4/DR5 (72–74), were significantly downregulated in both R77Q- and WT-infected cells compared to mock. The downregulation was more pronounced and significant in the R77Q infection (Fig. S2). This suggests a restrained extrinsic apoptotic pathway via a TRAIL-DR4/DR5 pathway in both infections (72). In addition, there was a coordinated downregulation across multiple TNF-family components such as *TNFRSF10D, TNFRSF12A,* and *TNFRSF10B* indicating dampened extrinsic apoptotic and inflammatory responses, consistent with viral modulation aimed at limiting caspase 8-dependent cell death (72, 75).

In the intrinsic apoptotic pathway, across both infections, the DEG findings support a strong intrinsic apoptotic pressure, but we noted a key difference in how effectively each viral strain engages the anti-apoptotic *bcl-2* family of genes. Prior work has shown that Vpr can directly and indirectly interact with the mitochondria and promotes intrinsic, caspase-independent apoptosis through mitochondrial permeabilization and Bcl-2 downregulation (39, 76). Consistent with that framework, our DEGs at 72 hpi point to a stressed, intrinsic apoptotic condition in both R77Q and WT infections, yet with different survival responses.

In WT-infected cells, anti-apoptotic gene expression is preserved. Compared to R77Q, WT infection shows higher expression of multiple anti-apoptotic factors in the *bcl-2* family including *bcl-2*, *bcl2l1/bcl-xl* and *mcl1*. This pattern fits the broader HIV-1 literature in which infected cells can simultaneously experience pro-death signals while maintaining survival programs that delay mitochondrial outer membrane permeabilization (MOP), supporting continued viral production or necrotic cell death (77, 78). R77Q, by contrast, shows a failure to induce this anti-apoptotic response. The most striking counterweight is the selective significant increase in the BH3-only activator *bcl2l11* (*bim*), a potent antagonist of *bcl-2* that lowers the threshold for MOP (34). In this context, the R77Q phenotype can be interpreted as a loss of function phenotype rather than a uniquely stronger pro-apoptotic trigger. R77Q infected cells thus appear to be less able to mount the WT-associated *bcl-2* upregulation, allowing intrinsic apoptosis to proceed once mitochondrial stress accumulates. This adds a novel mechanistic layer to prior observations that the apoptotic phenotype of Vpr is highly context- and sequence-dependent. When taken together, our data argues that both viruses engage intrinsic apoptotic stress, but the WT virus better preserves mitochondrial integrity via *bcl-2* upregulation, whereas R77Q lacks this compensatory response, shifting the *bcl-2/BH3* balance toward execution. This framework aligns with growing interest in leveraging cell-death pathways as therapeutic strategies in HIV (79), including efforts to target Bcl-2 family dependencies (79, 80). Notably, the Bcl-2 inhibitor Venetoclax has been explored as a means to sensitize HIV-1–infected cells to apoptosis by blocking Bcl-2–mediated survival (80, 81). Furthermore, our PPI analysis reinforces the idea that R77Q induces apoptosis by weakening the pro-survival interaction architecture rather than simply amplifying pro-apoptotic signals.

As with any transcriptomic study, our data does not fully capture post-transcriptional regulation (64), therefore, we envision that in the future, protein-level and mechanistic experiments will be important to fully capture the host response to the R77Q mutant. Importantly, the strengths of this approach lie in providing an unbiased, genome-wide view of how a single Vpr polymorphism reshapes host-cell fate, which is an essential foundation before dissecting downstream functional steps. Although *in vitro* systems cannot reproduce every aspect of *in vivo* infection, they offer the controlled environment necessary to reveal the distinct apoptotic mechanism induced by the R77Q mutant. Taken together, these design choices provide a rigorous and interpretable framework for understanding how the R77Q mutant influences apoptotic signaling and HIV-1 pathogenesis, setting the stage for more targeted mechanistic studies.

## Conclusion

In summary, these data support a model in which both viruses impose intrinsic apoptotic stress, but only WT infection elicits an anti-apoptotic response that prevents apoptosis. R77Q lacks this mechanism, making infected cells more susceptible to apoptosis once stress accumulates. The novel element of our findings is the demonstration that R77Q’s apoptotic phenotype arises not from a uniquely stronger pro-death signal, but from a loss-of-function phenotype that prevents the upregulation of key anti-apoptotic genes. This mechanistic distinction provides a plausible explanation for the reduced inflammatory profile linked to R77Q infection and offers insight into its association with slower disease progression.

## Materials and Methods

### Cell lines and cell culture

HEK293T cells were maintained at 37^°^C, 5% CO_2_ in DMEM (Sigma-Aldrich, USA), supplemented with 10% fetal bovine serum (Sigma-Aldrich, USA) and 1% penicillin/streptomycin (Sigma-Aldrich, USA). Ghost R3/X4/R5 cells were maintained at 37^°^C, 5% CO_2_ in DMEM, supplemented with 10% FBS, 1% P/S, 500 µg/mL G418, 100 µg/mL hygromycin, and 1 µg/mL puromycin. Ghost cells were obtained through the NIH HIV Reagent Program, Division of AIDS, NIAID, NIH: GHOST (3) CCR3+ CXCR4+ CCR5+ Cells from Drs. Vineet N. Kewal Ramani and Dan R. Littman (cat# 3934). HUT78 cells were maintained at 37 ℃, 5% CO_2_ in RPMI (Sigma-Aldrich, USA), supplemented with 10% FBS and 1% penicillin/streptomycin.

### Plasmids

HIV-1 pNL4-3 was obtained through the NIH HIV Reagent Program, Division of AIDS, NIAID, NIH: Human Immunodeficiency Virus 1 (HIV-1), Strain NL4-3 Infectious Molecular Clone (pNL4-3), ARP-114, contributed by Dr. M. Martin. The HIV-1 NL4-3 R77Q Vpr mutant was a gift from Dr. Velpandi Ayyavoo (University of Pittsburgh).

### Virus production and titration

Virus stocks were generated by transfection of HEK293T cells using the calcium phosphate method. Supernatants were harvested 48 hours post-transfection. Viral titers were quantified in Ghost R3/X4/R5 cells following the assay protocol provided by the NIH HIV Reagent Program (Rockville, MD, USA).

### HIV-1 infection assays

Infections were performed in 1 × 10^5^ HUT78 cells at an MOI of 0.3 in the presence of 10 µg/mL polybrene. Mock-infected cells were treated in the same concentration of polybrene and an equivalent volume of virus-free supernatant. All conditions were performed in triplicate.

### RNA Extraction and Sequencing

Total RNA was extracted from each condition at each time point using the Direct-zol^TM^ RNA MicroPrep w/TriReagent (Zymo Research, USA) extraction kit according to the manufacturer’s instructions. RNA purity and concentration were initially measured using the Thermo Scientific NanoDrop before being shipped to LC Sciences, Houston, TX, for poly-dT based mRNA library preparation and sequencing. RNA integrity was then assessed using the Agilent Technologies 2100 Bioanalyzer (High Sensitivity DNA Chip). All samples demonstrated high RNA integrity, with RNA Integrity Numbers (RIN) ≥ 8.6 and the majority exceeding 9.0, indicating suitability for downstream sequencing analysis. Sequencing was performed on a Illumina NovaSeq 6000 following their internal protocol and the generated FASTQ files were shared via a LC Sciences server. The sequencing reads and processed data obtained during this study have been submitted to the Gene Expression Omnibus (GEO) database, under accession number GSE319139.

### ARMOR and Quality Control

Downloaded FASTQ files were run through an existing Automated Reproducible Modular workflow for processing and differential analysis of RNA-seq data (ARMOR) v1.5.10 pipeline as previously described (82, 83). The ARMOR workflow performed quality control on the sequencing reads (FastQC v0.11.9), trimming (TrimGalore! v0.6.6) of adapters and poor-quality regions and mapping (Salmon v1.4.0) and quantifying sequencing reads to the human transcriptome. This workflow was performed on the MaryLou supercomputer at Brigham Young University, Provo, Utah, USA, using default parameters. The data processing steps are outlined in Fig. 8 as a summary.

**Fig. 8.**
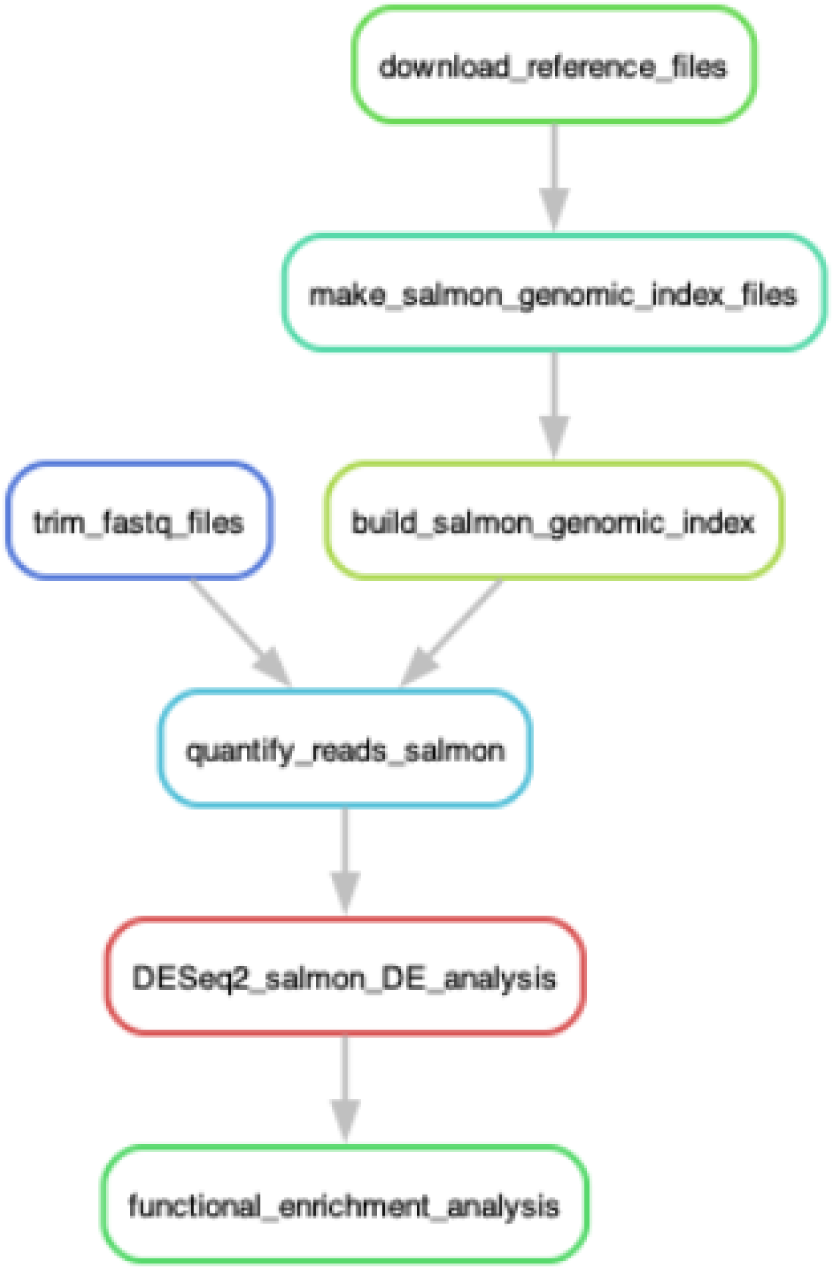
RNA seq data processing flow. Downloaded FASTQ files were processed using the Automated Reproducible Modular workflow for RNA-seq data (ARMOR, v1.5.10). The pipeline included read quality assessment (FastQC v0.11.9), adapter and low-quality base trimming (TrimGalore! v0.6.6), and transcript-level alignment and quantification against the human transcriptome (Salmon v1.4.0). Differential expression (DE) analysis was carried out using DESeq2 package v1.50.2.

### DEGs and functional enrichment analysis

Differential expression (DE) analysis was carried out using DESeq2 package v1.50.2 (84) which employs a negative binominal distribution model for determining statistical significance. Raw gene-level counts were imported, filtered to remove low-abundance features, and normalized using the trimmed mean of M-values (TMM) method (85, 86). Genes with a false discovery rate (FDR) adjusted p-value below 0.05 were considered significant.

Differential gene expression was followed by functional enrichment analysis using clusterProfiler v4.18.4 and gprofiler2 v0.2.4. GO term relationships and gene–term connections were visualized using enrichplot v1.30.3, which maps significant DEGs to hierarchical GO structures and highlights shared gene membership across terms (87, 88). To evaluate apoptosis-related responses, we filtered enriched GO biological process terms for keywords associated with apoptotic regulation (“apoptosis,” “programmed cell death,” and “cell death regulation”). Over-representation testing was conducted using a hypergeometric distribution and the Benjamini–Hochberg multiple hypothesis testing correction procedure, and pathway-level visualizations were generated with Pathview v1.41.1 (58).

### STRING protein-protein interaction

Predicted protein-protein interaction networks were generated using Search Tool for the Retrieval of Interacting Genes/Proteins (STRING) (v12.0; https://string-db.org) by submitting the list of significantly differentially expressed genes from the R77Q vs. WT comparison at 72 hpi. Network reconstruction was performed using the default settings, including the full STRING evidence channels (89). The resulting network was exported in both image and tabular formats, the latter containing interaction edges with associated confidence scores. Network visualization and refinement were carried out in the STRING interface, with node size scaled according to degree centrality and edge thickness mapped to interaction confidence to improve interpretability and figure quality.

### Statistical analysis

Differential gene expression was evaluated using standard linear modeling with false-discovery rate (FDR) p-adjusted correction (Benjamini–Hochberg). Genes with a FDR < 0.05 were considered significant.

## Data availability

The RNA sequencing reads were deposited in the Gene Expression Omnibus (GEO) Sequence Read Archive of the National Center for Biotechnology Information (http://www.ncbi.nlm.nih.gov/geo/) under accession no. GSE319139.

## Acknowledgements

We would like to thank the BYU Office of Computing for their assistance and access to the supercomputer as we analyzed our RNA sequencing data.

---

**Fig. S1. KEGG Cell Cycle pathway enrichment in R77Q vs. WT at 72 hpi**. The KEGG cell cycle pathway was visualized using Pathview, with differential gene expression data from the R77Q vs. WT comparison at 72 hpi overlaid onto the pathway map. Genes significantly upregulated in R77Q relative to WT are shown in red, while significantly downregulated genes are shown in blue. Uncolored nodes indicate genes that were not differentially expressed. The predominance of red nodes reflects broad upregulation of cell cycle–associated genes in R77Q compared to WT infection

**Fig. S2. TNF signaling pathway analysis in R77Q vs. WT at 72 hpi.** The KEGG TNF signaling pathway was visualized using Pathview, with differential gene expression data from the R77Q vs. WT comparison at 72 hpi overlaid onto the pathway map. Genes significantly upregulated in R77Q relative to WT are shown in red, whereas significantly downregulated genes are shown in blue. Uncolored nodes represent genes that were not differentially expressed. The predominance of blue nodes indicates broad downregulation of TNF signaling components in R77Q compared to WT infection.

**Supplementary table S1:** Top 10 most significant DEGs filtered in the R77Q vs. WT comparison.

| Gene name | Log2FC | FDR | General Related Function |
| --- | --- | --- | --- |
| DUSP5 | -2.01 | 1.87E-10 | Apoptosis |
| ICAM1 | -1.18 | 4.62E-10 | Inflammation |
| UNG | 1.08 | 9.97E-09 | Inflammation |
| CDCP1 | -2.12 | 2.48E-08 | DNA damage repair (BER) |
| PLD6 | 1.07 | 4.24E-08 | Mitochondrial fusion |
| ULBP2 | -1.16 | 1.22E-07 | Inflammation |
| ARG2 | -1.25 | 1.22E-07 | Mitochondrial protein |
| SERPINB9 | -1.41 | 1.70E-07 | Apoptosis |
| TREML2 | -1.23 | 1.86E-07 | Inflammation |
| SUPT16H | 0.714 | 2.21E-07 | DNA replication and repair |

